# DNAscent v2: Detecting Replication Forks in Nanopore Sequencing Data with Deep Learning

**DOI:** 10.1101/2020.11.04.368225

**Authors:** Michael A. Boemo

## Abstract

The detection of base analogues in Oxford Nanopore Technologies (ONT) sequencing reads has become a promising new method for the high-throughput measurement of DNA replication dynamics with single-molecule resolution. This paper introduces DNAscent v2, software that uses a residual neural network to achieve fast, accurate detection of the thymidine analogue BrdU with single-base resolution. DNAscent v2 comes equipped with an autoencoder that detects replication forks, origins, and termination sites in ONT sequencing reads from both synchronous and asynchronous cell populations, outcompeting previous versions and other tools across different experimental protocols. DNAscent v2 is open-source and available at https://github.com/MBoemo/DNAscent.

---

The high-throughput detection of replication fork movement with single-molecule resolution is critical for understanding how a cell replicates its DNA and how this can be exploited for theraputic purposes. Oxford Nanopore Technologies (ONT) sequencing has emerged as a cost-effective platform for the detection of DNA base modifications such as 5-methylcytosine on long single molecules [1–5]. We and others have shown that halogenated bases are also detectable in ONT sequencing data [6–9]. When these bases are pulsed into S-phase cells, they are incorporated into nascent DNA by replication forks (Figure 1a). Sequencing with ONT and detecting the position of these bases reveals a footprint of replication fork movement on each sequenced molecule, allowing this method to answer questions that would have been traditionally addressed with DNA fibre analysis but with higher-throughput and the ability to map each sequenced read to the genome. DNAscent (v1 and earlier) uses a hidden Markov model to assign a likelihood of BrdU to each thymidine [7], RepNano uses a convolutional neural network to estimate the fraction of thymidines substituted for BrdU in rolling 96-bp windows [8], and NanoMod compares modified and unmodified DNA to detect base analogues [5, 6]. As this method matures, it is critical that software is able to detect base analogues with high accuracy and throughput across different experimental protocols in a way that is easy to use. To that end, this paper introduces DNAscent v2 which uses a residual neural network to accurately assign a probability of BrdU to each thymidine and an autoencoder to detect replication forks, origins, and termination sites at any point in S-phase. This work demonstrates that DNAscent v2 is the new state-of-the-art to support DNA replication and genome stability research.

**Figure 1:**
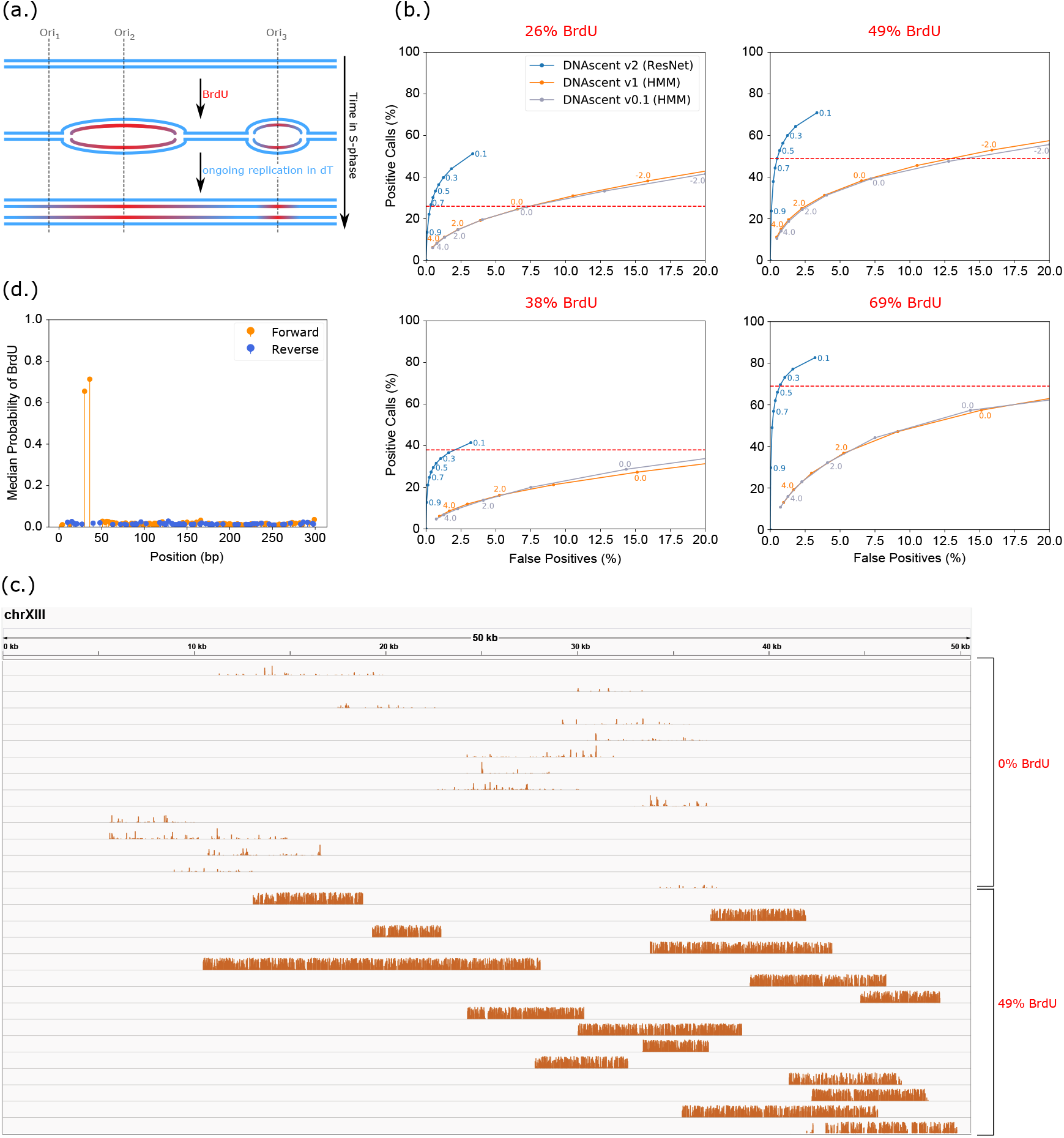
Performance of the DNAscent v2 `detect` subprogram. **(a)** When the thymidine analogue BrdU is pulsed into S-phase cells, BrdU is incorporated into the newly replicated nascent DNA in place of thymidine. Detecting BrdU in nascent DNA sequenced with ONT can reveal the movement of replication forks in millions of single molecules. **(b)** ROC curves showing the ratio of positive BrdU calls to false positive BrdU calls for four different experiments with different BrdU-for-thymidine substitution rates. The 26% and 49% BrdU samples are from [7] while the 38% and 69% samples are from [8]. The BrdU-for-thymidine substitution rate as measured by mass spectrometry is indicated by the dashed red line. Points along each curve are different thresholds above which a BrdU call is considered positive; for DNAscent v2, these are probabilities whereas for DNAscent v1 and below, these are log-likelihoods. Each curve was calculated using 5,000 reads. The x-axis of each plot has been truncated from 0-100% to 0-20% for clarity. **(c)** Bedgraphs visualised in IGV [10] showing the proability of BrdU called at each thymidine position for a randomly selected subset of reads used in the 49% BrdU ROC curve analysis. Each track is a single read, and the y-axis of each track ranges from 0 to 1. **(d)** The median probability of BrdU called by DNAscent v2 at each thymidine position along primer extension reads (N=273) from [7] where BrdU has been substituted for thymidine in two known positions (30 and 36 bp) on the forward strand. All other positions on the forward and reverse strand are unsubstituted.

The subprogram `detect` in DNAscent v2 detects BrdU with single-base resolution by using a residual neural network consisting of depthwise and pointwise convolutions (Figure S1). The model was trained using nanopore-sequenced genomic DNA from a *S. cerevisiae* thymidine auxotroph [7]. In particular, the training material consisted of unsubstituted DNA as well DNA with 80% BrdU-for-thymidine substitution (see Section S1 for details). To determine the ratio of true positives to false positives, receiver operator characteristic (ROC) curves were plotted using nanopore sequenced unsubstituted DNA to measure false positives and DNA with four different BrdU-for-thymidine substitution rates (Figure 1b). DNAscent v2 outperformed previous versions by a wide margin in all four samples. Bedgraphs of the probability of BrdU at each thymidine position for a subset of unsubstituted reads and 49% BrdU-for-thymidine substituted reads from the ROC curve analysis are shown in Figure 1c, highlighting the difference between substituted and unsustituted reads. In concordance with the ROC curves, unsubstituted reads are largely devoid of false positives. To show that DNAscent v2 distinguishes BrdU from thymidine with single-base resolution, BrdU detection was run on substrates with two BrdU bases at known positions [7] where DNAscent v2 was able to clearly identify the positions of both BrdU bases (Figure 1d).

DNAscent v2 includes a new subprogram called `forkSense` that uses an autoencoder neural network to assign the probabilities that a leftward- and rightward-moving fork passed through each position on a read during the BrdU pulse (see Section S2 for details). `forkSense` matches up converging and diverging forks in order to call replication origins and termination sites on each nanopore-sequenced molecule. A shortcoming of earlier DNAscent versions was that origin calling was designed to work in synchronised early S-phase cells, but `forkSense` works in synchronous and asynchronous cells at any point in S-phase. DNAscent v2 was tested on two different BrdU-pulse experimental protocols: *S. cerevisiae* cells that were synchronised in G1 and released into S-phase in the presence of BrdU with no thymidine chase [7] and asynchronous thymidine-auxotrophic *S. cerevisiae* cells where BrdU was pulsed for 4 minutes followed by a thymidine chase [8]. Example single molecules mapping to a region that includes several efficient origins on *S. cerevisiae* chromosome I are shown for both experiments (Figure 2a-b). With DNAscent v2, the BrdU calls are clean enough that the single-base resolution BrdU calls can be visualised directly as bedgraphs in IGV [10] without the need for any smoothing or further processing from the software. `forkSense` calls origins as the regions between diverging leftward- and rightward-moving forks and calls termination sites as the regions between converging forks. A pileup of replication origins and termination sites called on *S. cerevisiae* chromosome II is shown for cells synchronised in G1 (Figure 2c) and asynchronous cells (Figure 2d). While the location of called replication origins shows good agreement with confirmed and likely origins from OriDB [11] in both cases (Figure 2e-f) this work corroborates the findings of [7, 8] that high-throughput, single-molecule analysis reveals replication origins that are far (> 5 kb) away from previously annotated origins. DNAscent v2 is able to capitalise on its improved BrdU detection to detect several fold more origins than both previous versions of DNAscent and RepNano (Figure 2e-f).

**Figure 2:**
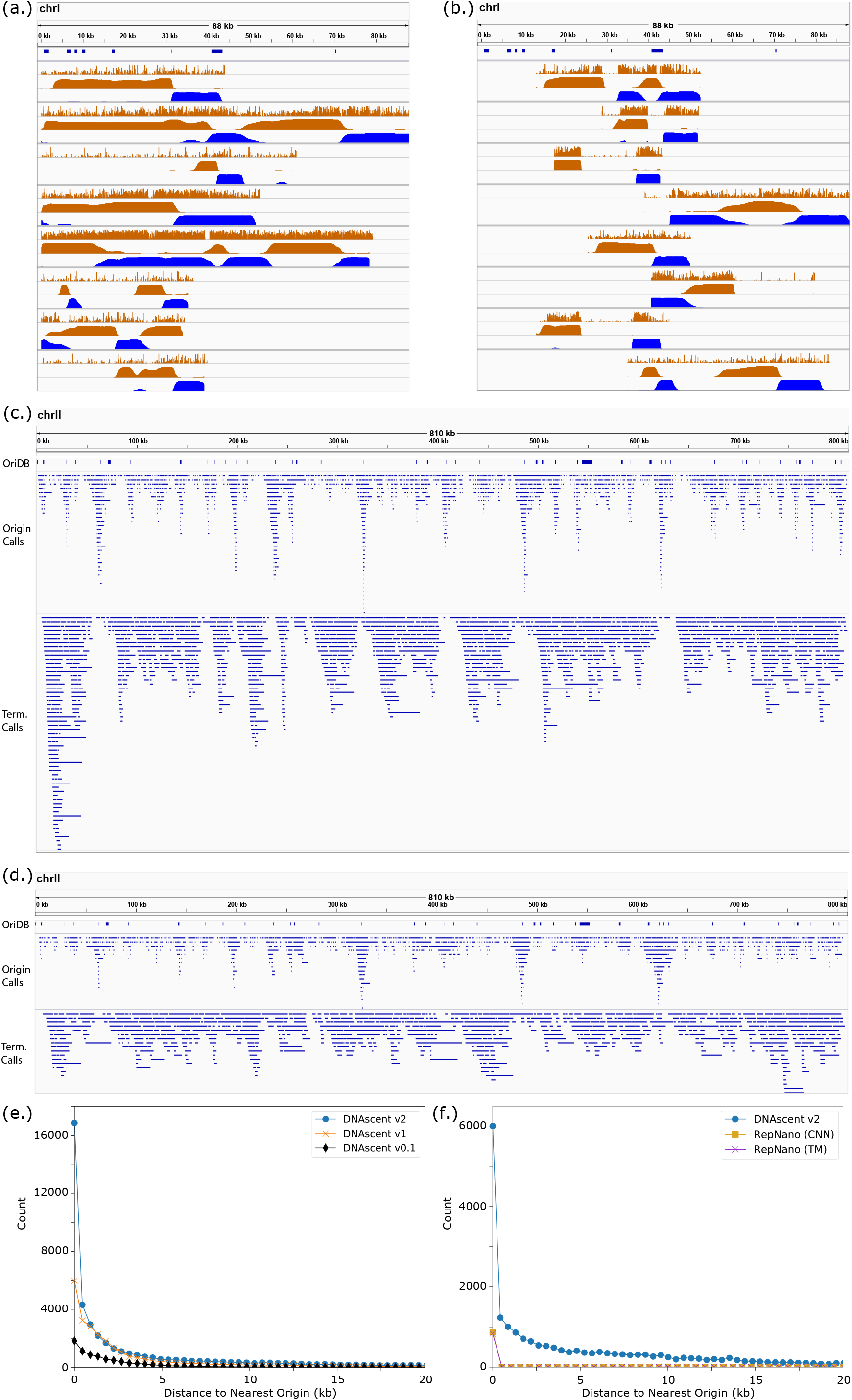
Performance of the DNAscent v2 `forkSense` subprogram. **(a-b)** Individual reads mapping to *S. cerevisiae* chromosome I are shown for *S. cerevisiae* cells **(a)** synchronised in G1 and released into BrdU and **(b)** asynchronous and pulsed with BrdU and chased with thymidine. Origins that are confirmed and likely from OriDB are shown in the top track. Eight reads are shown for each experiment where each read is represented as a group of three tracks: the probability of BrdU at each thymidine (upper track; from DNAscent detect) and the probability that a leftward-moving fork (middle track; from DNAscent `forkSense`) and rightward-moving fork (lower track; from DNAscent `forkSense`) passed through each position during the BrdU pulse. The y-axis of each track ranges from 0 to 1. **(c-d)** Pileup of all replication origins and termination sites called by `forkSense` that mapped to *S. cerevisiae* chromosome II for *S. cerevisiae* cells **(c)** synchronised in G1 and released into BrdU (2,980 origin calls from a total of 9,864 reads) and **(d)** asynchronous and pulsed with BrdU and chased with thymidine (1,461 origin calls from a total of 5,186 reads). Only reads with mapping length ≥ 20 kb and mapping quality ≥ 20 were used. The OriDB track shows confirmed and likely origins. **(e)** Distribution of the distance between each origin call and the nearest confirmed or likely origin for *S. cerevisiae* cells synchronised in G1 and released into BrdU. The results of three versions of DNAscent are shown. RepNano only made a total of 14 origin calls on this dataset when run with the default settings, so these were omitted for clarity. **(f)** A similar analysis to Figure 2e, but for asynchronous *S. cerevisiae* cells pulsed with BrdU and chased with thymidine. Results for DNAscent v2 are shown alongside results from the RepNano transition matrix (TM) and convolutional neural network (CNN) origin calling algorithms run using the default parameters. Earlier versions of DNAscent were not designed to call origins in asynchronous cells, so only the results from DNAscent v2 are shown.

Transitioning the DNAscent `detect` BrdU calling algorithm from the hidden Markov forward algorithm to residual neural networks has advantages beyond significantly improving the ratio of true positives to false positives; it serves as a more natural platform for future work on the detection of multiple base analogues in the same molecule. DNA fibre analysis relies on sequential pulses of different base analogues to determine fork direction while DNAscent currently determines fork direction from the changing frequency of BrdU-for-thymidine substitution across a molecule. while DNAscent’s current approach is advantageous in its simplicity, the detection of multiple analogues would be necessary to answer certain questions typically addressed with fibre analysis, such as the stability of stalled replication forks [12]. The improvement to single-base resolution BrdU calling in `detect`, together with the `forkSense` algorithm, has allowed DNAscent v2 to make significantly more origin calls than previous versions when run on the same dataset, and as shown by Fig. 2e, most of these additional calls were near confirmed and likely origin sites. This suggests a decrease in false negative origin calls, enabling DNAscent v2 to create a more accurate picture of how replication took place on each individual molecule. With these improvements, less data is required to create whole-genome maps of replication origin and termination site locations, which is particularly important for studying replication in larger genomes.

In addition to improving performance and adding functionality, DNAscent v2 development placed a particular focus on ease-of-use and accessibility for laboratories that may not have access to computational scientists or bioinformaticians. Origin calling with RepNano has fourteen adjustable parameters and earlier versions of DNAscent have three, but `forkSense` in DNAscent v2 does not require any tuning. DNAscent v2 also comes packaged with a utility that converts the outputs of `detect` and `forkSense` into bedgraphs such that BrdU and fork probabilities can easily be viewed side-by-side for each read (as in Figure 2a-b) in IGV [10] or the UCSC Genome Browser (http://genome.ucsc.edu) [13], and origin, termination, and fork calls are likewise written to bed files. To support the analysis of larger datasets, DNAscent v2 can optionally run BrdU detection on a GPU and benchmarks approximately 4.5× faster than DNAscent v1 and approximately faster than RepNano (Section S3).

This paper has introduced DNAscent v2, which utilises residual neural networks to drastically improve the single-base accuracy of BrdU calling compared with the hidden Markov approach utilised in earlier versions. DNAscent v2 also includes the new `forkSense` subprogram which uses an autoencoder to infer the movement of replication forks from patterns of BrdU incorporation. `forkSense` can call the location of replication forks, origins, and termination sites in single-molecules across a range of experimental protocols with a sensitivity that exceeds both earlier versions and other competing tools. These new methodologies, together with improvements in speed and ease-of-use, make this technology an important new piece of the toolkit in DNA replication and genome stability research.

## Data Access

DNAscent v2 is open-source under GPL-3.0 and is available at https://github.com/MBoemo/DNAscent. ONT sequencing data for BrdU detection training, primer extension, and synchronised cell cycle experiments were released with [7] in NCBI GEO under accession number GSE121941. ONT sequencing data for the asynchronous cell cycle experiment was released with [8] in ENA under accession number PRJEB36782 (experiment ERX4016778).

## Acknowledgements

The author would like to thank Dr. Carolin Müller, Dr. Rosemary Wilson, and Dr. James Carrington (Sir William Dunn School of Pathology, University of Oxford), Dr. Conrad Nieduszynski (Earlham Institute), Dr. Mathew Jones (Diamantina Institute, University of Queensland), Dr. Jared Simpson (Ontario Institute for Cancer Research and University of Toronto), as well as Dr. Catherine Merrick and Dr. Francis Totanes (Department of Pathology, University of Cambridge) for helpful conversations and critical reads of this manuscript.

