## Supplemental Information for "DNAscent v2: Detecting Replication Forks in Nanopore Sequencing Data with Deep Learning"

### Supplementary Information

Michael A. Boemo*[1]

[1]Department of Pathology, University of Cambridge, Tennis Court Road, Cambridge CB2 1QP, United Kingdom

#### S1 BrdU Detection Residual Neural Network and Training

The architecture of the residual neural network used by `detect` for single-base BrdU detection is shown in Figure S1. The model is comprised of five blocks of depthwise and pointwise one-dimensional convolutions followed by batch normalisation with hyperbolic tangent activation, and the model architecture is loosely based on [1]. The input of each block is subject to pointwise convolution and batch normalisation, where it is then added to the end of the block. All convolutional layers within the same block have the same kernel size and number of filters, and these values are specified for each block in Table S1.

To create the model's input, Oxford Nanopore reads were basecalled with Guppy (version 3.3.3) and aligned to the reference genome using minimap2 (version 2.17-r941). BrdU incorporation leads to significant inaccuracies in Guppy's basecalling, so the ground-truth sequence for each read was taken to be the subsequence of the reference genome that the read aligns to. The raw Oxford Nanopore signal was segmented into events using Scrappie (https://github.com/nanoporetech/scrappie) and aligned to this ground-truth sequence using an adaptive banded alignment [2] in order to normalise for shift and scale. Following normalisation, a more precise hidden Markov event alignment was done using methods similar to nanopolish `eventalign` (https://github.com/jts/nanopolish) [3]. Using the hidden Markov alignment, the read was transformed into an $8 \times M$ tensor where $M$ is the number of unique positions on the reference genome the read matched to during the hidden Markov event alignment. The eight features for each reference position are the one-hot encoded alphabet $\{A, T, G, C\}$ for the base at that reference position, the mean of the events aligning to that position, the sum of the event lengths (in seconds) of all events aligning to that position, as well as the expected Oxford Nanopore event mean and standard deviation for the sequence at that position. The output of the softmax layer is a $2 \times M$ dimensional tensor specifying the probability the position is a BrdU and the probability that the base is in $\{A, T, G, C\}$ for each position on the reference that the read aligns to.

The model was trained on a total of 75,000 reads from [4] of which 50,000 were unsubstituted and 25,000 had 80% BrdU-for-thymidine substitution as measured by mass spectrometry. To account for uncertainty in which positions were BrdU in the 80% substituted reads, fuzzy labels were assigned to each position via bootstrapping with the hidden Markov model BrdU detection algorithm in DNAscent v1. The DNAscent v1 `detect` algorithm was run on the 25,000 BrdU-substituted reads and positions with a log-likelihood of BrdU greater than 1.25 were taken to be a positive call. Positions called were given the label,

$$P(\text{position is BrdU} \mid \text{positive HMM call}) = \frac{TP \cdot C}{TP \cdot C + FP \cdot (1 - C)}$$

and positions called as thymidine were given the label,

$$P(\text{position is BrdU} \mid \text{negative HMM call}) = \frac{FN \cdot C}{FN \cdot C + TN \cdot (1 - C)}$$

where $C$ is the fraction of BrdU-for-thymidine substitution measured by mass spectrometry and $TP$, $FP$, $FN$, and $TN$ are the true positive, false positive, false negative, and true negative rates for HMM-based BrdU detection in DNAscent v1, respectively. The true positive rate used for HMM-based detection was 0.5 and the true negative rate used was 0.9. Both of these values are lower than the results in Figure 1b (orange curve) suggest are appropriate, but testing determined that conservative estimates of these parameters led to better training results. While the 25,000 BrdU-substituted reads were labelled according to HMM-based BrdU detection bootstrapping, the 50,000 unsubstituted reads were given uniform label smoothing of $P(\text{position is BrdU}) = 0.01$ for all thymidine positions.

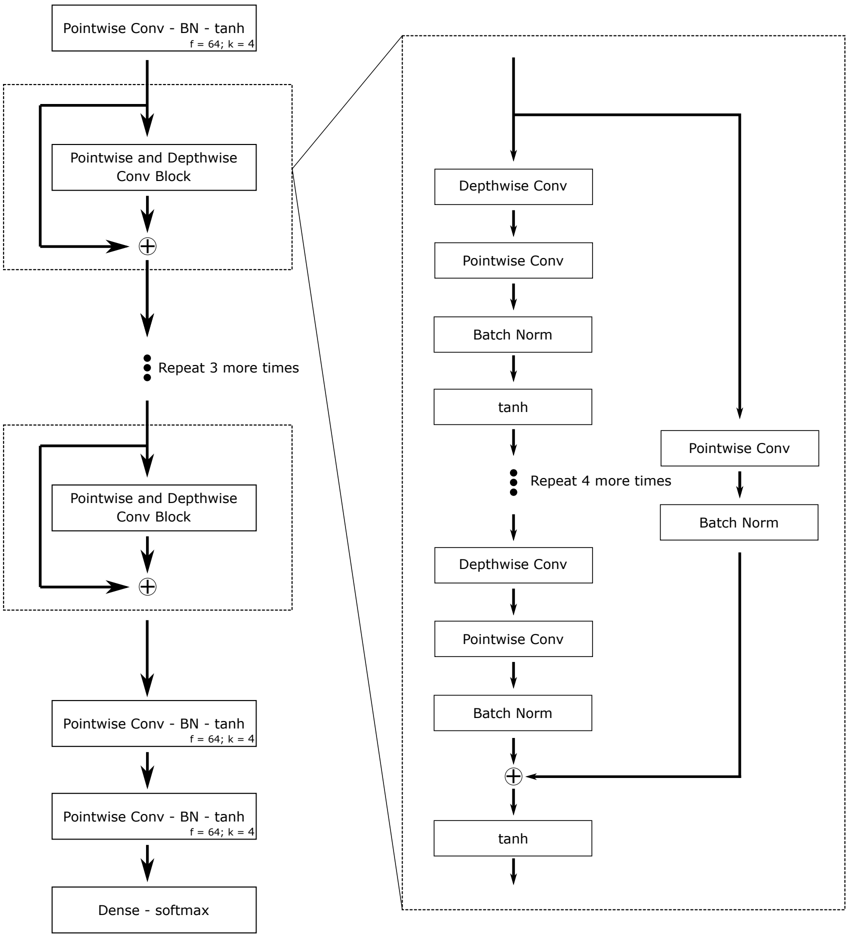


Figure S1: Architecture of the residual neural network used in single-base BrdU detection. The number of filters ($f$) and the kernel size ($k$) for convolutional layers are specified. The filters, kernel size, and strides for each block (dotted lines) are specified by Table S1. The total number of parameters in the model is 1,512,962.

Data augmentation in the training set was done as shown in Figure S2. When a BrdU-substituted read and an unsubstituted read covered the same region of the reference genome, if a position on the substituted read was called as BrdU by HMM-based detection, the events from the substituted and unsubstituted read aligning to a 6 bp window around that position were swapped. The input tensor for these augmented reads were built as described above, and the labels for each position in the augmented read were unchanged from the read the events were taken from.

| Block | Filters | Kernel Size | Stride |
|---|---|---|---|
| 1 | 64 | 4 | 1 |
| 2 | 64 | 4 | 1 |
| 3 | 128 | 8 | 1 |
| 4 | 128 | 8 | 1 |
| 5 | 256 | 16 | 1 |

Table S1: The number of filters, kernel size, and strides of each of convolutional layer (both pointwise and depthwise) for each the five blocks in the residual neural network outlined in Figure S1.

The model was implemented and trained using Tensorflow (version 1.14.0). Training was done using an adaptive moment estimation optimizer with categorical crossentropy loss. The model was trained with batches of 32 reads for a total of 9 epochs.

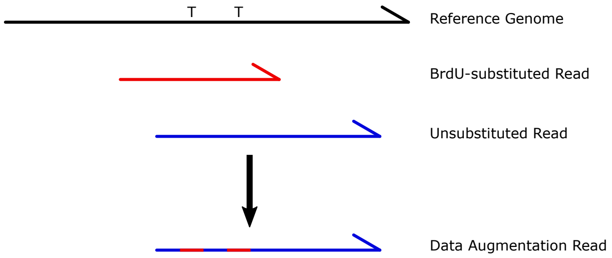


Figure S2: Data augmentation strategy for DNAscent v2 `detect` training. A BrdU-substituted read (red) and an unsubstituted read (blue) both map to the same region of the genome. Where the primary sequence of the two reads overlap, if the two thymidines indicated were called as BrdU by DNAscent v1 `detect`, then the events aligning to a 6 bp window around these thymidines are swapped between the BrdU-substituted read and unsubstituted read to make a new augmented read. These augmented reads help by establishing a more reliable ground truth across the read of which positions are BrdU and which are thymidine.

### S2 Fork Direction Autoencoder and Training

The architecture of the `forkSense` autoencoder that calls replication forks from the output of DNAscent `detect` is shown in Figure S3. The model consists of pointwise convolutions and max pooling layers (the encoding) followed by pointwise convolutions and upsampling (the decoding). The details of each convolution, pooling, and upsampling layer are given in Table S3. The input to the model is a $1 \times M$ tensor created from the output of `detect` which consists of the probability of BrdU at each of the $M$ positions on the read that align to the reference genome. The model outputs a $3 \times M$ tensor that, for each position, specifies the probability that a rightward-moving fork moved through that position during the pulse, the probability that a leftward-moving fork moved through that position during the pulse, and the probability that no fork moved through that position during the pulse.

The model was trained on 1027 reads from [4] that showed high BrdU incorporation around known origins of replication in *S. cerevisiae* and 500 "blank" reads from [4] that had no BrdU incorporation. Note that these training reads were excluded from the analysis in Figure 2. For each of the 1027 reads with BrdU incorporated around a known origin, the BrdU calls were smoothed with a moving-average filter and leftward- and rightward-moving forks were labelled as starting from the origin and ending at the point where the smoothed BrdU probabilities dropped below 0.1, with all other positions labelled as "no fork". To account for inaccuracies in labelling and prevent over-confident calls, label smoothing was applied to all positions according to

$$L_{smooth} = L(1-\alpha) + \frac{\alpha}{K}$$

where $L = 1$ is the unsmoothed label for the class, the model distinguishes between $K = 3$ classes, and using the label smoothing parameter $\alpha = 0.06$ [5].

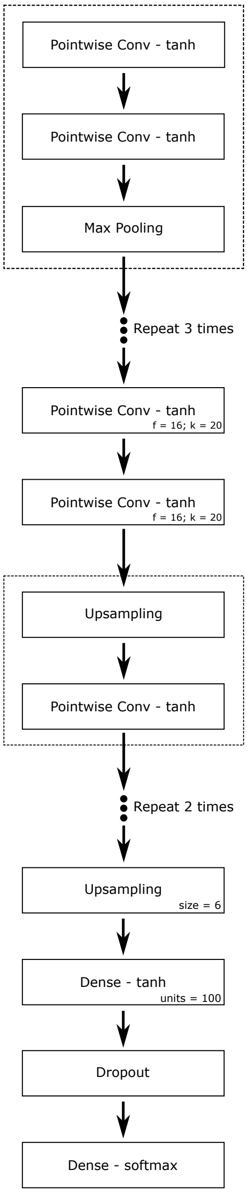


Figure S3: Architecture of the autoencoder neural network used to detect replication forks from the output of DNAscent `detect`. The number of filters ($f$) and the kernel size ($k$) for convolutional layers are specified. The filters, kernel size, and strides for each block (dotted lines) are specified by Table S2. The total number of parameters in the model is 16,271.

| Block | Filters | Kernel Size | Stride | Pooling/Upsampling Size | Pooling Stride |
|---|---|---|---|---|---|
| 1 | 4 | 10 | 1 | 10 | 6 |
| 2 | 4 | 13 | 1 | 10 | 4 |
| 3 | 8 | 15 | 1 | 10 | 4 |
| 4 | 8 | 17 | 1 | 10 | 4 |
| 5 | 8 | 17 | 1 | 4 | - |
| 6 | 4 | 15 | 1 | 4 | - |
| 7 | 4 | 10 | 1 | 4 | - |

Table S2: The number of filters, kernel size, and strides of each of convolutional layer (both pointwise and depthwise) for each the seven blocks in the autoencoder outlined in Figure S3.

Data augmentation was done for each of the 1027 BrdU-substituted reads as shown in Figure S4. Each read was divided at the origin in order to create four additional augmented reads: an isolated rightward-moving fork, an isolated leftward-moving fork, forks converging in a termination site, and separated forks consistent with an origin that fired before the start of the BrdU pulse. This scheme carried a number of advantages. First, it enabled training on only one run and one experimental protocol so that others could be used as proper test set. Second, as the cells were synchronised with BrdU pulsed at the start of S-phase, both forks were guaranteed to be near the origin. This prevented having to make *a priori* assumptions about whether patterns of BrdU observed away from cannonical origins should be labelled as isolated leftward-moving forks, isolated rightward-moving forks, termination sites, or replication initiation that had occured away from known origins.

As in the previous section, the `forkSense` model was implemented and trained using Tensorflow (version 1.14.0) and training was done using an adaptive moment estimation optimizer with categorical crossentropy loss. The model was trained with batches of 32 reads for a total of 25 epochs.

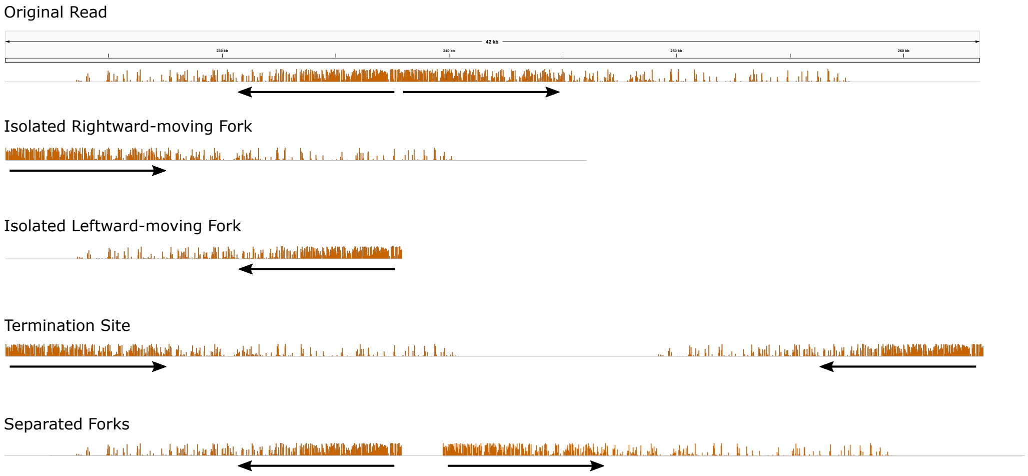


Figure S4: Data augmentation strategy for DNAscent v2 `forkSense` training. Each training read was split at the known origin of replication to create reads with isolated forks, forks converging in a termination site, and separated diverging forks.

### S3 Runtime

BrdU detection for all three tagged DNAscent releases (v0.1, v1, and v2) and RepNano (commit fd129c1; 5 May 2020) were benchmarked on a single bulk fast5 file consisting of 4000 reads from *S. cerevisiae* cells synchronised in G1 and released into BrdU [4]. Reads in this file were basecalled with Guppy (version 3.3.3). All benchmarked software was run three times on these reads using 12 CPUs (Intel® Xeon® Gold 6142 Skylake @ 2.60GHz). DNAscent v2 has the added functionality of running BrdU prediction on a GPU, so DNAscent v2 was also run using 12 CPUs (Intel® Xeon® CPU E5-2650 v4 Broadwell @ 2.20GHz) and an NVIDIA® Tesla® P100 16 GB GPU. The runtime for each trial is shown in Table S3.

| | Trial 1 | Trial 2 | Trial 3 | Average |
|---|---|---|---|---|
| DNAscent `detect` v2 (GPU) | 00:14:55 | 00:14:37 | 00:14:30 | 00:14:41 |
| DNAscent `detect` v2 (CPU) | 00:19:21 | 00:18:55 | 00:18:53 | 00:19:03 |
| DNAscent `detect` v1 | 01:10:22 | 01:05:35 | 01:06:30 | 01:07:29 |
| DNAscent `detect` v0.1 | 00:28:13 | 00:26:35 | 00:26:29 | 00:27:06 |
| minimap2 alignment | 00:00:21 | 00:00:20 | 00:00:18 | 00:00:20 |
| RepNano `preprocess` | 00:07:05 | 00:03:49 | 00:06:06 | 00:05:40 |
| RepNano `predict_simple` | 00:47:38 | 00:48:17 | 00:48:12 | 00:48:02 |

Table S3: Runtimes for BrdU detection with DNAscent `detect` and RepNano on one bulk fast5 file of 4000 reads. All times are in the format hours:minutes:seconds.

To benchmark the total runtime of the BrdU detection pipeline for each software tool, the starting point was taken to be Guppy-basecalled reads and the end point was taken to be BrdU calls across those reads. For RepNano, this was the summed runtimes of `preprocess` and `predict_simple`. It is important to note that RepNano does a minimap2 alignment internally; to compare with DNAscent, the runtime of a minimap2 alignment was summed with the runtime of DNAscent `detect`. The runtime of each pipeline is shown in Table S4.

| | Pipeline Runtime | Relative Runtime |
|---|---|---|
| DNAscent `detect` v2 (GPU) | 00:15:01 | 1.00× |
| DNAscent `detect` v2 (CPU) | 00:19:23 | 1.29× |
| DNAscent `detect` v1 | 01:07:49 | 4.52× |
| DNAscent `detect` v0.1 | 00:27:26 | 1.82× |
| RepNano | 00:53:42 | 3.57× |

Table S4: Runtime of BrdU detection pipelines using different software tools. All times are in the format hours:minutes:seconds.

Unless the sequencing run is very large, running `forkSense` on the output of `detect` is not a major contributor of compute time to the workflow. Table S5 shows the runtime of `forkSense` (with replication fork, origin, and termination calling enabled) when run on the output of `detect` for the same 4,000 reads used in Tables S3 - S4. `forkSense` was run three times on the same dataset using 1 CPU and 12 CPUs (Intel® Xeon® Gold 6142 Skylake @ 2.60GHz).

| | Trial 1 | Trial 2 | Trial 3 | Average |
|---|---|---|---|---|
| DNAscent `forkSense` (12 cores) | 00:00:23 | 00:00:18 | 00:00:17 | 00:00:19 |
| DNAscent `forkSense` (1 core) | 00:00:40 | 00:00:39 | 00:00:40 | 00:00:40 |

Table S5: Runtime of DNAscent `forkSense` on one bulk fast5 file of 4000 reads. All times are in the format hours:minutes:seconds.
